# Changes in the microbial community of *Lubomirskia baicalensis* affected by Brown Rot Disease

**DOI:** 10.1101/641811

**Authors:** ROREX Colin, BELIKOV Sergej, BELKOVA Natalia, Chernogor Lubov, Khanaev Igor, Nalian Armen, Martynova-Van Kley Alexandra

## Abstract

Sponge diseases occur globally and the resulting reduction of sponge populations has negative effects on other organisms within the ecosystems due to loss of nutrient enrichment and loss of bioremediation. In Lake Baikal, the predominate sponge species Lubomirskia baicalensis is currently being infected with an unidentified pathogen resulting in a sharp decline in population. The current hypothesis is that the recent increase in methane concentration in the lake has caused dysbiosis within the bacterial community of L. baicalensis resulting in the disease outbreak. In this study we investigated the changes in the bacterial community between healthy and sick sponges using 16S bacterial profiling targeting veritable regions 3-5. Here we present data that the bacterial communities of the healthy sponge samples were significantly different from sick samples and several poorly classified organisms were identified by Indicator Species Analysis as significant. Organisms identified from the sick samples classified within taxonomic units that contain acidophilic bacteria which suggest pH may play a role. There was also an observed decrease in the number of identified methyltropic bacteria present in the sick sponge samples compared to the healthy.

## Introduction

Lake Baikal in Siberia is the largest rift lake in the world and home to an estimated 20% of the world’s fresh water. It was formed 25 million years ago via plate tectonics and is a unique ecosystem that is of significant cultural, economic, and scientific value. Native to Lake Baikal are several species of freshwater sponge, of which *Lubomirskia baicalensis* is the predominate species. Currently, researchers at Lake Baikal are reporting an outbreak of disease affecting *L. baicalensis* [1 and 2].

This disease outbreak first occurred in 2011 [1]. Prior to this report there had been no historical record of systemic sponge disease in this location. The long-term effects of a severe sponge disease are difficult to predict. Losses in sponge population can result in increases of phytoplankton blooms with negative environmental and economic effects [3]. In addition, the loss of sponge mass can also negatively impact oxygen levels and nutrient availability in the ecosystem as sponges play critical roles as filter feeders, both consuming a wide range of microorganisms and increasing nutrient availability [4].

Sponge disease mechanisms can vary with some disease outbreaks caused by a single organism and others requiring multiple organisms. The disease outbreak in the Great Barrier Reef of *Rhopaloeides odorabile* was reported to have been caused by a novel α-proteobacteria [5]. In the Red Sea, microbiota recovered from diseased sponge tissues taken from two sites 30 km apart were both found to be colonized by the same species of verrucomicrobia [6].

Several outbreaks have been shown to require exposure to multiple pathogenic microorganisms. In a disease outbreak off the coast of Papua New Guinea, five bacterial species were isolated from diseased tissue and used to inoculate healthy sponge tissue. Here the disease state was only replicated in culture when five bacterial isolates were used together as an inoculum [7]. In Sponge Necrosis Syndrome, two bacteria and four fungal species were identified in the diseased tissue. Reproducing the disease in culture required a mixture of one bacterial and one fungal species [8].

The Sponge White Patch disease, originally thought to be the cause of a sponge boring bacteria infecting *Amphimedon compressa*, was re-evaluated utilizing bacterial microbiome profiling techniques. It was found that diseased sponges had a different microbiota than healthy sponges, with some of the species detected previously implicated in other sponge and coral diseases [9].

In the case of the sponge disease reported in Lake Baikal, the causative organism(s) or factors have not been identified. One hypothesis is that the disease is caused by a dysbiosis in *L. baicalensis* as a result of increasing methane concentrations in Lake Baikal [1].

This pilot study classifies the bacterial communities using five tissue samples collected before and after the disease outbreak occurred. Our hypothesis is that sick sponges will have a distinct bacterial community not shared by healthy sponges and that significant taxa occurring within the sick sponges may correlate to factors related to disease state.

## Materials and Methods

### Samples

Three samples of DNA extracted from sponge tissue were received from our collaborators at the Lake Baikal, Irkutsk, Limnological Institute SB RAS. Two samples were collected from healthy sponges (PI and PII), the third sample (PIII) was collected from a diseased sponge. Already sequenced data was provided for three additional samples. These additional samples were sequenced using the Roche 454 platform and include a duplicate of the diseased sample PIII-454, an uninfected sample Healthy-454, and a laboratory cultivated aggregates from dissociated single sponge cells called primmorphs (Table 1).

**Table 1.**
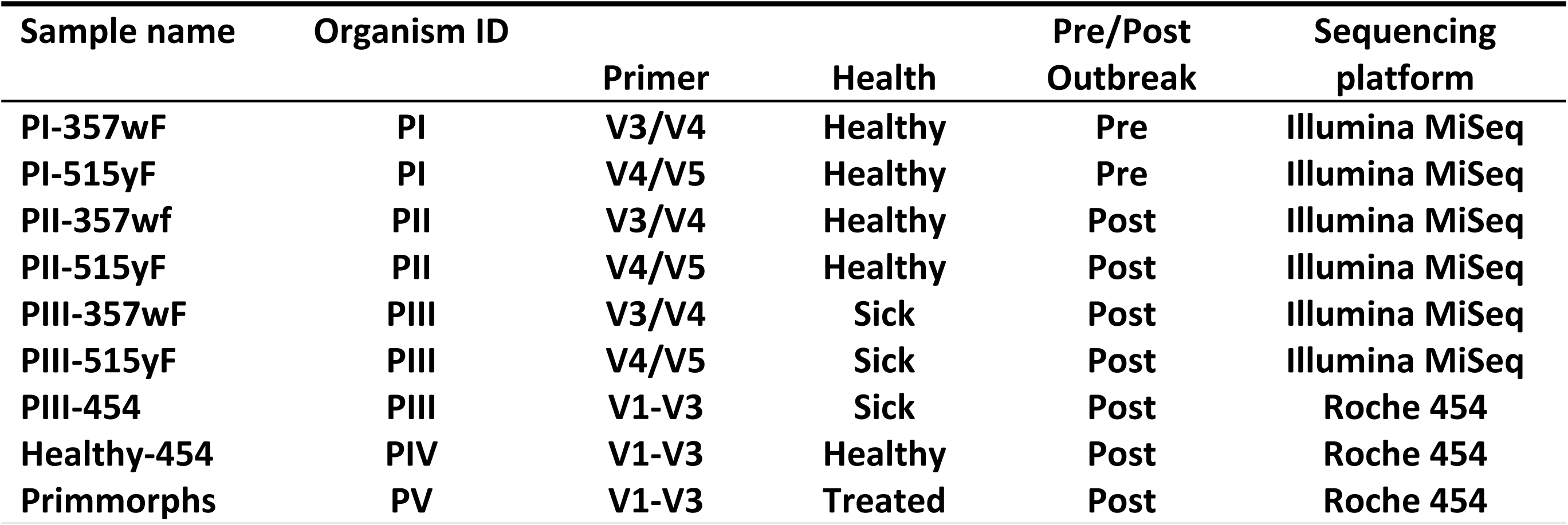
Description of all samples in this study. Sample name is the identifier for this study. Organism ID is an arbitrary label used when the same DNA pool is shared or when both samples are taken from the same organism. The primer refers to the 16S variable regions used to sequence the sample. Outbreak status indicates whether the sample was collected before or after the first signs of disease were detected. The sequencing platform indicates which NGS sequencing platforms was used to generate sequencing data.

| Sample name | Organism ID | Primer | Health | Pre/Post<br>Outbreak | Sequencing<br>platform |
| --- | --- | --- | --- | --- | --- |
| PI-357wF | PI | V3/V4 | Healthy | Pre | Illumina MiSeq |
| PI-515yF | PI | V4/V5 | Healthy | Pre | Illumina MiSeq |
| PII-357wf | PII | V3/V4 | Healthy | Post | Illumina MiSeq |
| PII-515yF | PII | V4/V5 | Healthy | Post | Illumina MiSeq |
| PIII-357wF | PIII | V3/V4 | Sick | Post | Illumina MiSeq |
| PIII-515yF | PIII | V4/V5 | Sick | Post | Illumina MiSeq |
| PIII-454 | PIII | V1-V3 | Sick | Post | Roche 454 |
| Healthy-454 | PIV | V1-V3 | Healthy | Post | Roche 454 |
| Primmorphs | PV | V1-V3 | Treated | Post | Roche 454 |

### Amplification and Sequencing

Amplification and sequencing of amplicons was conducted by RTL Genomics of Lubbock, Texas (www.rtlgenomics.com). Samples were amplified using two 16S rRNA primer sets: 357wF/785R for variable regions 3 and 4 (V3-V4) and 515yF/916yR for variable regions 4 and 5 (V4-V5). Sequencing was conducted on an Illumina platform.

### Primer sequences

357wF (V3-V4)

**CCTACGGGNGGCWGCAG**

785R (V3-V4)

**GACTACHVGGGTATCTAATCC**

515yF (V4-V5)

**GTGYCAGCMGCCGCGGTAA**

926pfR (V4-V5)

**CCGYCAATTYMTTTRAGTTT**

### Data Analysis

FASTA data was prepared by removing barcode information from sequence reads prior to taxonomic determination. Taxonomic data was generated for both Illumina and Roche 454 data sets using RDP classifier (rdp.cme.msu.edu) with an 80% confidence threshold. Both data sets were then merged into a unified matrix file. An environmental matrix was constructed with descriptive values for the samples such as primer, sequencing platform, disease condition, and collection date. The vegan package for R-studio (rstudio.com) was used for statistical analysis. Species data was visualized using the envfit function of vegan to construct a non-metric multidimensional scaling (NMDS) ordination and then fitted with descriptive variables to the ordination using 100,000 permutations. Indicator species analysis (ISA) was performed and the samples were organized into three higher order groups based on primer and clustering in ordination space. PI-515yF and PII-515yF were clustered into the Healthy 1 group. The Healthy 2 group consisted of PI-357wF, PII-357wF and Healthy-454. The Sick group consisted of the three PIII samples consisting of PIII-357wF, PIII-515yF and PIII-454. The primmorph sample was omitted from ISA as it contained only a single set of data.

## Results

### Species Richness

Species richness, the number of unique taxonomic groups in each data set, ranged from 37 taxonomic groups to 89 taxonomic groups. Samples amplified with the 515yF primer had the lowest species richness with 37 unique taxa with the PI sample, 40 with PII and 61 with the PIII sample. The samples amplified with the 357wF primer recovered more taxonomic groups with both PII and PIII recovering 77 unique taxa and PI 88. The Roche 454 samples, recovered 74, 78 and 89 for the PIII-454, Healthy-454 and primmorph samples respectively (Figure 1).

**Figure 1.**
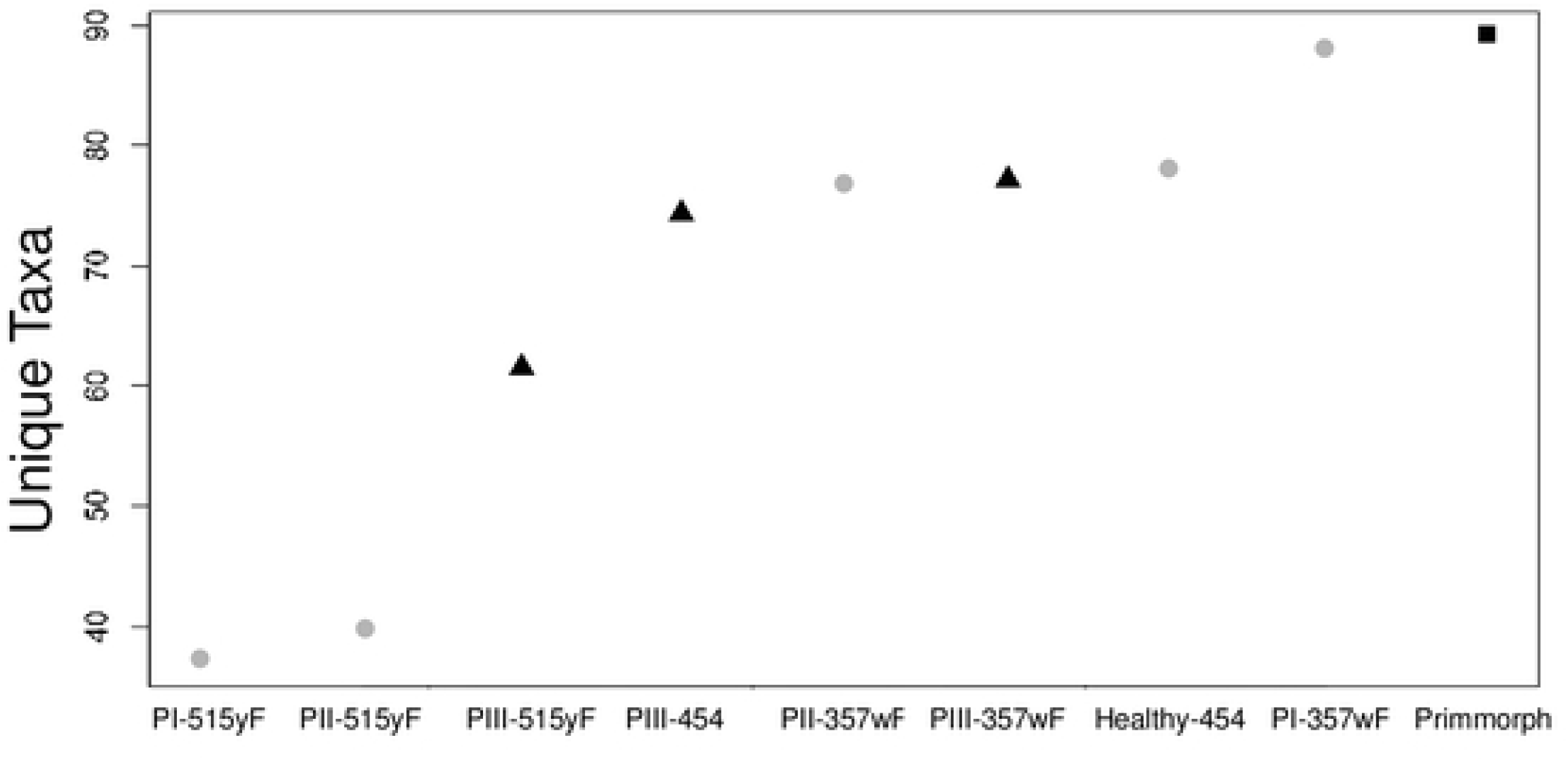
Species richness for all microbiome profiles. PI, PII and Healthy are samples obtained from uninfected sponge tissue. PIII are samples from an infected sponge. Primmorph is a sample that was cultured under laboratory conditions obtained from sponge tissue. Suffixes indicate primer set used 515yF for V4-V5 amplification, 357wF for V3-V4 or for samples sequenced on the Roche 454 (454) platform.

### NMDS

Multiple different environmental variables were fitted to the NMDS data to test for relationships between the samples in ordination space. A p-value of 0.4374 was obtained when attempting to fit the sequencing platform to the data, indicating that the underlying relationship structuring the points in ordination space is unrelated to the type of sequencer used. Likewise, attempting to fit primer data to the ordination failed to achieve statistical significance with a p-value of 0.4385. Other factors considered, such as collection year, had no relevance to the ordination structure (p-value 0.7981).

A significant p-value of 0.023 was obtained when the data was fitted against the disease status of the sponge samples (Figure 2). The three samples from the diseased sponge PIII-357wF, PIII-515yF and PIII-454 clustered together in ordination space. The five samples from health sponges, while showing a wide distribution in ordination space, still formed a cluster around the Healthy-454 sample. The primmorph sample did not group with either of the two other groups.

**Figure 2.**
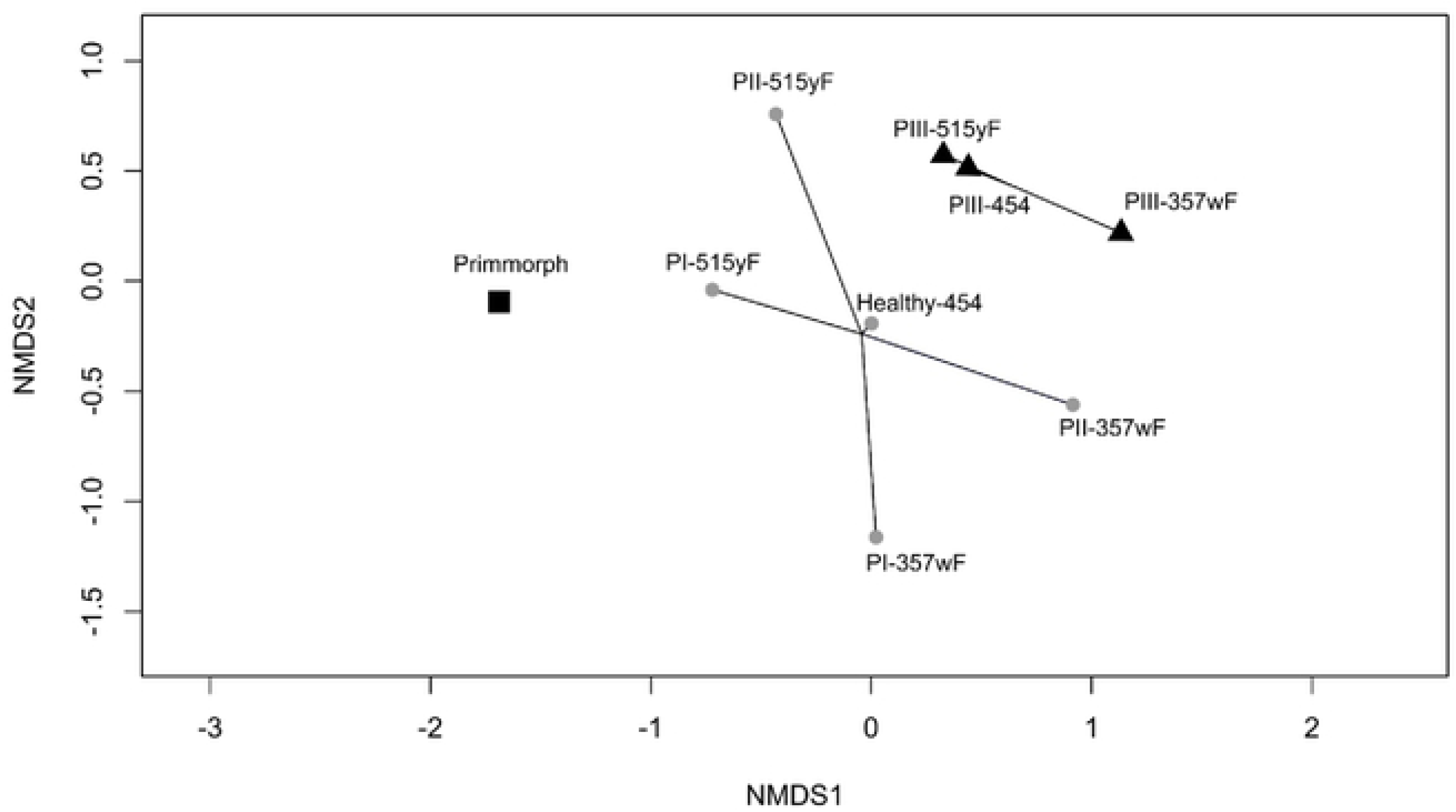
NMDS fitting sponge disease status. Healthy samples are represented by circles. Sick samples by triangles and the primmorph sample by a square. Samples are linked by health status (sick, healthy, primmorph). PI and PII (both 515yF and 357wF primers) and Healthy-454 cluster in one group. The diseased samples PIII (515yF, 357wF and 454) cluster. And the single primmorph sample is distinct from either groups.

### Indicator Species Analysis

Several significant taxa were identified as significant by ISA (Table 2). Only statistically significant samples (p < 0.05) were reported (Table 2). The Healthy 1 group, samples amplified with the V4-V5 primer, had no significant OTUs. Healthy 2 included an organism in the order Flavobacteriales, a broad classification of aquatic bacteria, and an organism in the family Clostridiaceae 1, which contains organisms of very different pathogenicities and ecological niches.

**Table 2.** Summary of the significant indicator species from all data sets. The sick group is a combined group of all PIII samples. The Healthy 2 group consists of the PI-357wF, PII-357wF and Healthy-454 samples. The group Healthy 1 did not contain any samples with a p < 0.05. A is the proportion of times a taxonomic group occurred within that group. B is the frequency the taxonomic group occurred in the samples that make up that group.

| Healthy 2 |  |  |  |  |
| --- | --- | --- | --- | --- |
|  | A | B | Stat | P value |
| Clostridiaceae 1 | 1 | 1 | 1 | 0.0396 |
| Flavobacteriales | 1 | 1 | 1 | 0.0396 |
| Sick |  |  |  |  |
|  | A | B | Stat | P value |
| Opitutus | 1 | 1 | 1 | 0.0344 |
| Acidobacteria Gp3 | 0.9778 | 1 | 0.989 | 0.023 |
| Acetobacteraceae | 0.84 | 1 | 0.916 | 0.0428 |
| Sick and Healthy 2 |  |  |  |  |
|  | A | B | Stat | P value |
| Actinobacteria | 1 | 1 | 1 | 0.0368 |
| Alcaligenaceae | 1 | 1 | 1 | 0.0368 |
| Candidatus pelagibacter | 1 | 1 | 1 | 0.0368 |

Three taxa were identified within the Sick group. Opitutus is a poorly understood genus of Verrucomicrobia found in rice paddy soil [10]. Acetobacteraceae is a family of oxidative fermenters that can tolerate low acid environments. Acidobacteria Gp3 is a subdivision of the phylum Acidobacteria which also contains known acidophiles.

The combined Healthy 2 and Sick group had several significant taxa common to aquatic ecosystems. *Pelagibacter unqiue* is the most abundant marine and freshwater bacterium on Earth [11]. Alcaligenaceae is a family of bacteria found in all non-extreme environments, some of which are known to be pathogenic. Terrimicrobium, a Verrucomicrobia, is another poorly-characterized bacteria that was first found in rice paddies. One organism classified only at the phylum level, Actinobacteria, has members which are ubiquitous in terrestrial and aquatic ecosystems.

### Methylotrophic bacteria

Only the V3-V4 primer set recovered sequence data that identified methylotrophic bacteria. The highest number of detected methylotroph sequences was in the PI samples, collected prior to the disease outbreak. We observed a decrease in the PII and PIII samples both collected after the disease outbreak with the lowest abundance in the sick PIII samples (Table 3).

**Table 3.** A table of **methylotrophic** organisms and the number of sequences recovered in each sample. Samples highlighted in gray were collected from sick organisms. Samples are separated by primer set.

|  | Methylo-<br>coccaceae | Methylo-<br>cystaceae | Methylo-<br>cystis | Methylo-<br>philaceae | Methylo-<br>soma | Methylo-<br>tenera |
| --- | --- | --- | --- | --- | --- | --- |
| PI-515yF | 0 | 0 | 0 | 0 | 0 | 0 |
| PII-515yF | 0 | 0 | 0 | 0 | 0 | 0 |
| PIII-515yF | 0 | 0 | 0 | 0 | 0 | 0 |
| PI-357wF | 0 | 0 | 0 | 401 | 0 | 144 |
| PII-357wF | 0 | 2 | 12 | 0 | 16 | 0 |
| Healthy-454 | 0 | 0 | 22 | 5 | 5 | 22 |
| PIII-357wF | 0 | 1 | 17 | 18 | 0 | 0 |

## Discussion

Based on the species richness data, the 515yF primer set does not recover the same depth of taxonomic units as the 357wF primer set. This is most likely due to differences in the primers rather than a sample specific phenomenon. Both primer sets amplify the V4 region along with a different additional variable region. The choice of which variable region to sequence does have a noticeable impact on the type of data recovered [12]. One of the limitations of the 515yF-926pfR primer set was the lack of any recovered methyltroph sequences compared against the V1-V3 primer and the V4-V5 primer.

Multiple descriptive factors were tested against the NMDS to determine if they were likely to explain the distribution. Among the factors tested were sequencing platform, health status of the samples, primer used for amplification, *L. baikalensis* organism sampled, and whether the samples were collected before or after the outbreak. The only statistically significant factor was the health of the organism with a p value of 0.0243. This indicates the bacterial communities of the sick samples are distinct from those of the healthy samples. This also drove the creation of the higher order groups used in ISA.

ISA identified two organisms in taxonomic groups known to contain acidophiles, Acidobacteria gp3 and acetobacteraceae. Since these two organisms occur almost exclusively within the sick samples, 97.8% and 84% respectively, this suggests either these organisms or a low pH may be a correlating factor to disease.

Since methane and methyltrophs are hypothesized to be involved in the disease outbreak, it was unexpected that there was an observed decrease in abundance of methyltrophs in the sick samples. This suggests that methyltrophs could be a transient component of the sponge microbiome during the course of infection with initial colonization of methyltrophs causing a dysbiosis event that enables pathogenic microorganisms to colonize causing disease. Then the observed decrease in methyltrophs may result from the changing post-infection bacterial community where other microorganisms are able to outcompete the methyltrophs.

In conclusion, we find that the bacterial community of sick sponges is distinct from that of healthy sponges. The identification of two poorly-classified acidophiles significant to the sick samples should be further investigated. To fully explore this both a larger data set needs to be obtained and an alternate V4-V5 primer used as the 515yF-926pfR primer did not recover any sequences classifying as methyltrophs. We propose that in addition to the collection of more samples, pH measurements both in the immediate environment and in sponge tissue should be collected.

## Acknowledgements

This study was supported by the Government Contract no. VI.50.1.4, “Molecular Ecology and Evolution ….” (no. 0345-2015-0002).

